# Understanding RNA Chaperone Activity of ProQ Protein

**DOI:** 10.1101/2025.01.17.633555

**Authors:** Shilpi Singh, Nisha Kumari, Tanmay Dutta, Tarak Karmakar

## Abstract

RNA-binding proteins (RBPs) are crucial in recognizing and binding small RNAs that modulate gene expression. RBPs often act as RNA chaperones, facilitating the sRNA-mRNA duplex formation. One of the most studied sRNA chaperones is Hfq, the post-transcriptional gene regulator in bacteria. Recently, a new RNA-chaperone protein, ProQ, has been discovered, which binds to a novel sRNA *RaiZ*, derived from the 3’-end of mRNA encoding ribosome-inactivating protein, and promotes its base pairing with the ribosome binding site of hupA mRNA, leading to the repression of HU-α protein synthesis. Despite these studies, the RNA-binding mechanism in ProQ has not been well explored in detail to date. In this work, we studied the binding of *raiZ* RNA to ProQ using atomistic long-timescale molecular dynamics and on-the-fly probability-based enhanced sampling (OPES) simulations. The *raiZ* RNA binds to ProQ’s concave surface, mostly decorated with polar and positively charged residues like Arginine and Lysine, exerting electrostatic stabilization to the RNA. *In-silico* mutations of these crucial residues lead to a significant loss of the intermolecular interactions, resulting in the detachment of the RNA from the protein. The free energy surface obtained from the OPES simulation helps identify stable *raiZ-*bound ProQ structures as well as intermediate states. The pertinent atomistic insights into the protein-RNA interactions and deciphering the binding mechanism of the *raiZ* RNA to ProQ enrich our knowledge about ProQ and its RNA chaperone activity.

## 1. Introduction

RNA-protein interactions play a pivotal role in many physiological processes in all domains of life.^1^ The outcome of these interactions includes the transfer of genetic information from DNA to proteins and the regulation of gene expression circuits in the cell.^2,3^ Among the many RNAs, a class of regulatory RNAs in bacteria termed small RNAs (sRNAs) finetune the gene expression^4,5^ in the cell to help them adapt to changing environments.^6^ sRNAs primarily modulate the gene expression at the post-transcriptional level through base pairing to their cognate targets, and in many instances, require a chaperone protein. One of these extensively characterized matchmaker proteins that promote sRNA-mRNA base pairing is Hfq.^7^ However, recent studies using Grad-seq^8^, CLIP-seq^9^, and RIP-seq identified a new RNA-chaperone protein ProQ^10^, which was identified earlier as an osmoregulatory factor obligatory for the expression of proline transporter protein ProP in *Escherichia coli*^11^, its RNA-binding capability had never been examined.

The initial indication of the RNA binding activity of ProQ stemmed from the structural similarities of its N-terminal domain (NTD) with the crystal structure of an RNA binding protein (RBP), Fertility inhibitor O (*FinO*). Subsequent ProQ RNA interactome studies have established ProQ as an RBP^12^, which acts as a bacterial RNA chaperone^13^ and binds largely to a different set of *trans*-acting sRNAs that bind to Hfq.^14^ The higher affinity of ProQ for double-stranded RNA indicates that RNA with stem-loop structure(s) is a major determinant for ProQ binding to it. However, not all sRNAs with stem-loop structure(s) have been observed to bind ProQ. This points to additional structural specificities of sRNAs for ProQ binding. Investigations on the ProQ-bound sRNAs revealed that ProQ binds to the 3’-terminus of sRNAs comprising a prerequisite stem-loop followed by a single-stranded poly-Uracil (U) stretch. The 3’-end stem-loop of sRNA with less than two U residues is unable to bind to ProQ, which instigates an understanding of the mechanism of how ProQ interacts with sRNA and eventually promotes its binding to target mRNA.

ProQ comprises two globular domains connected through an elongated bridging intradomain flexible linker. The N-terminal FinO-like domain possesses RNA binding activity, while the C-terminal domain (CTD) consists of a Tudor domain fold, which acts to interpose protein-protein interactions.^15^ Although NTD of ProQ has been reported to bind independently to sRNAs, coordinated interactions from NTD, CTD, and linker domain of ProQ are necessary for an out-and-out binding to sRNA, as NTD alone is insufficient to bind some typical sRNAs. Knowledge of the mode of interaction of ProQ with sRNA ligands is inadequate as only a handful of studies have been conducted so far to understand the mechanism completely. ProQ has been shown to bind a novel sRNA, *raiZ* derived from the 3’-end of *raiA* mRNA encoding ribosome-inactivating protein, and promotes its base pairing with the ribosome binding site of *hupA* mRNA, leading to the repression of HU-α protein synthesis.^16^ Of the few other ProQ-sRNA interaction studies, the one that demonstrated that ProQ binds two different sRNA ligands, *rbsZ* and *rybB*, together and facilitates their base pairing interaction^17^, happens to follow a distinct mechanistic pathway. The lack of ample experimental studies to establish the ProQ binding mechanism to sRNAs drove us to investigate the mechanism of ligand binding to ProQ systematically.

In this work, we have investigated the binding mechanism of a small RNA *raiZ* to the ProQ N-Terminal (Figure 1a) using extensive atomistic molecular dynamics (MD) and on-the-fly probability-based enhanced sampling (OPES) simulations.^18^ The *raiZ* binds to the concave surface of the protein in a semi-folded state. Strong electrostatic interactions between the positively charged residues, such as ARG, HIS, LYS, and the backbone phosphate group of *raiZ*, help the guest bind in the ProQ pocket. The RNA binding mechanism and the thermodynamics of the binding process have been studied using OPES simulations using Deep-TDA collective variable (CV)^19^ that can constructively capture binding and unbinding events of ProQ-*raiZ*. Furthermore, we performed *in silico* mutations of important amino acids, ARG69, ARG80, ARG100, and LYS101 (Figure 1b); mutating them to ALA resulted in the unbinding of *raiZ* (Figure 1c) from the ProQ binding pocket.

**Figure 1:**
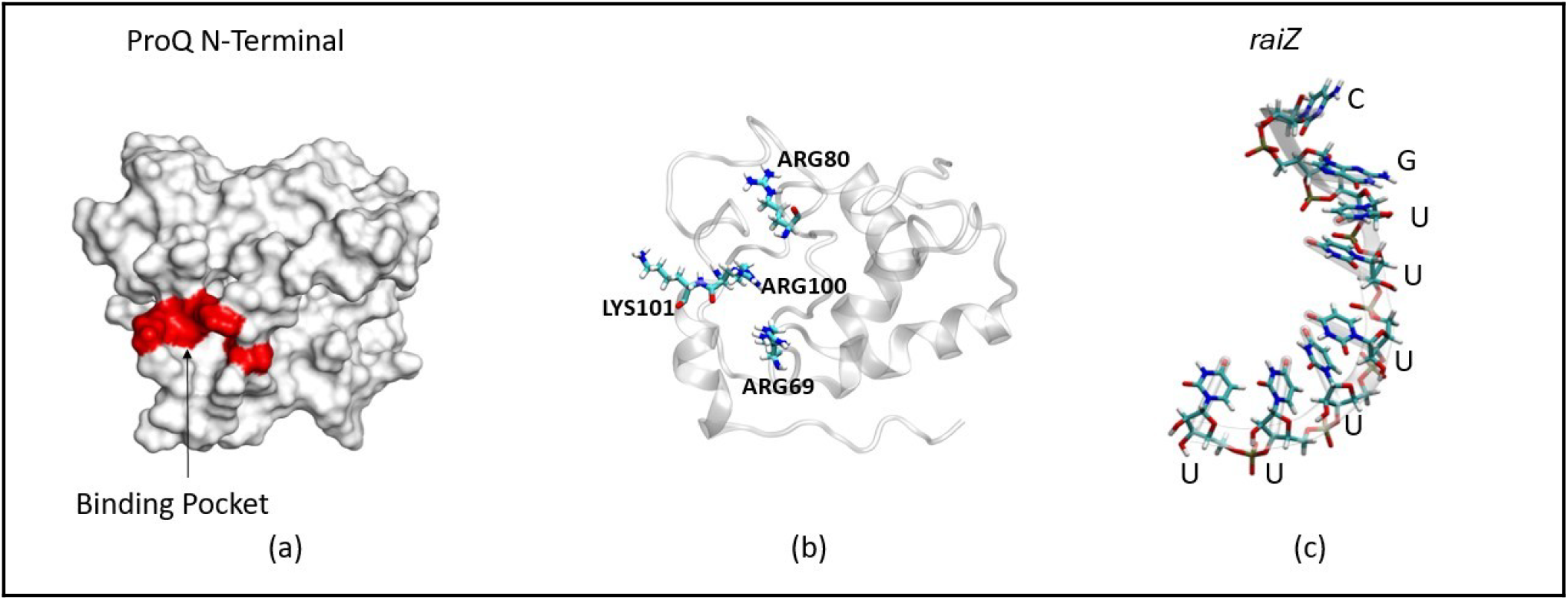
(a) ProQ N-Terminal (surface representation) (PDB ID: 5nb9), the red highlighted region shows the binding pocket of ProQ, (b) ProQ N-Terminal (cartoon representation) shows important ARG residues for binding *raiZ*, and (c) 3’ end of sRNA *raiZ* consists of 8 nucleotides 5’-CGUUUUUU-3’

## 2. Results & Discussion

### 2.1. Conformational flexibility of ProQ and *raiZ*

To understand the conformational behavior of ProQ, we carried out an equilibrium MD simulation of a single ProQ-NTD in water. Backbone root mean square deviations (RMSDs) show that ProQ retains its structural integrity throughout the simulation time (Figure S1-a in SI). The high value of RMSD is because of the flexible loops (residues 1-12 and 110-117) present in the ProQ structure. We subsequently conducted an MD simulation of an eight-nucleotide segment of *raiZ* small RNA (5’-CGUUUUUU-3’) to elucidate its conformational behavior in water. We selected the 8-nucleotide-long *raiZ* because its 3’ end contains more than two uracils, which are crucial for binding and are part of the stem loop. In Figure S1-b, the RMSD fluctuations are due to the free extended structure of *raiZ;* no secondary structure of *raiZ* has been observed to form during this simulation.

### 2.2 *raiZ* binding modes and the stability of ProQ-*raiZ* complex

Once we comprehended the individual behavior of ProQ and *raiZ* in water, our curiosity was heightened to investigate the behavior of the ProQ-*raiZ* complex. Unfortunately, the crystal structure of the *raiZ-*bound ProQ structure is not available. We docked the *raiZ* into ProQ using AutoDock Vina software^20^ and obtained a few representative structures of the ProQ-*raiZ* complex, each with varying binding energy values (Table S1 in SI). The structure with the highest binding energy was subjected to a 10 μs-long MD simulation to check the complex’s stability and conformational behavior.

From the MD simulation trajectory, we found that the *raiZ* is tightly attached to the concave surface of ProQ, resulting in the formation of a distinctive w-shaped structure of *raiZ* RNA (State-1) (Figure 2a). Our investigation examined the intermolecular interactions that stabilize the *raiZ* RNA on the ProQ surface. The RNA is stabilized on the ProQ surface by electrostatic interactions between the positively charged amino acid residues ARG32-U7, ARG69-U3, ARG109-C1, and ARG80-U8 and the *raiZ’s* negatively charged backbone (Figure S2-a in SI). In addition, the RNA in the ProQ-*raiZ* complex is further stabilized by hydrogen bonds formed between PRO76-U8, SER26-U6, GLN118-U5, TRP68-G2, and TYR78-U3 (Figure S2-b in SI).

**Figure 2:**
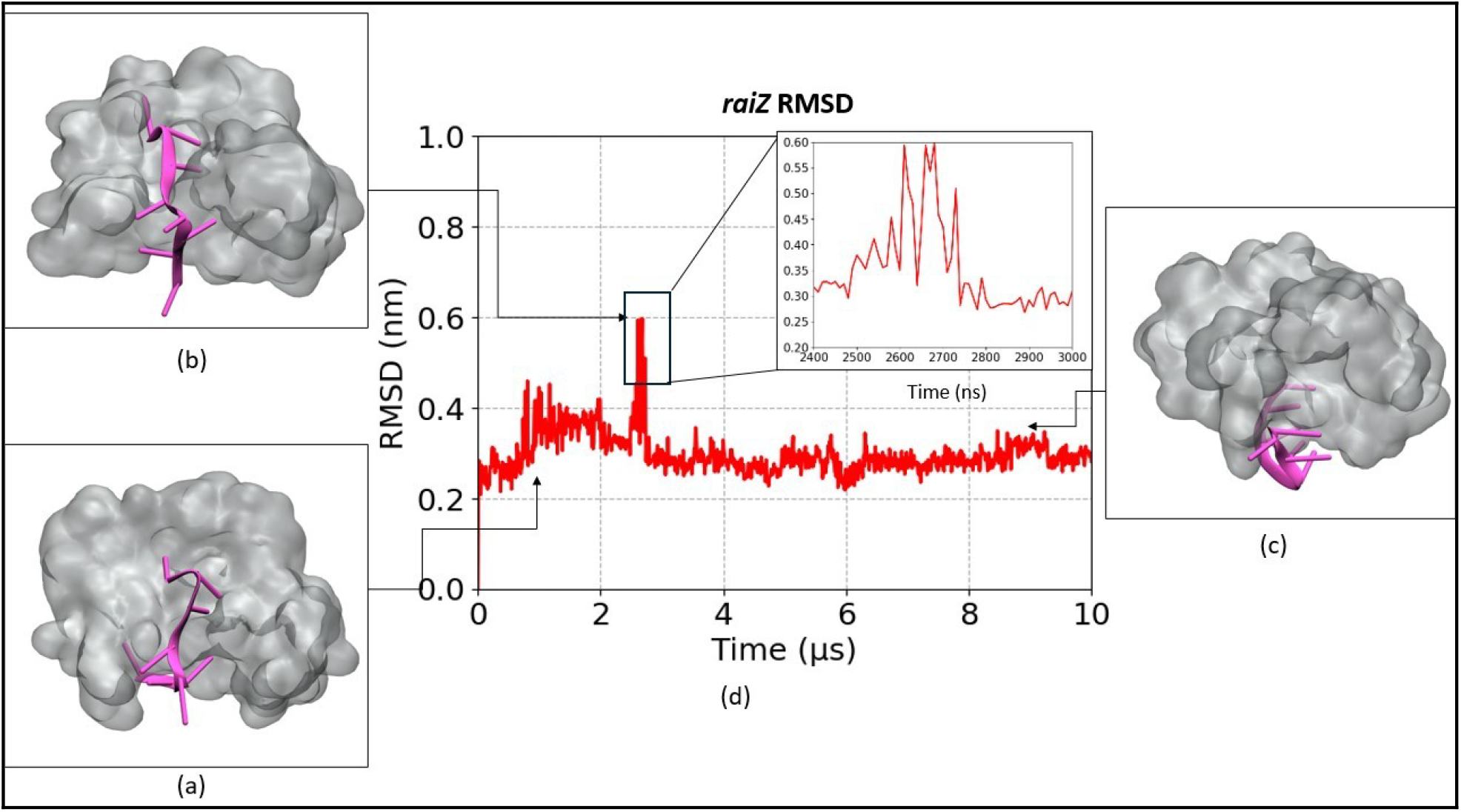
(a) Binding poses of *raiZ* inside ProQ pocket in State-1 (b) intermediate state shows switch from State-1 to State-2, (C) Binding poses of *raiZ* inside ProQ pocket in State-2 (d) RMSD of *raiZ* bound to ProQ

From the MD simulation trajectory, we observed that the *raiZ* changes its position after about 2.7 microseconds (as shown in Figure 2b).

These changes are accompanied by a notable rise in the *raiZ* backbone RMSD (Figure 2d). As a result of this change in the position of *raiZ*, the interactions in State-1 (Figure 2a) were disrupted, causing it to transition towards the second conformation state, State-2 (Figure 2c). In State 2, *raiZ* adopted a loop-shaped structure and attached it to the lower side of the ProQ protein. Specifically, the 3’ end of *raiZ* is bound to the lower backside residues of ProQ. As a result, new hydrogen bonds were formed between ProQ and *raiZ*. Electrostatic interactions occurred between positively charged amino acid residues and the RNA backbone involving PHE39-C1, SER59-U8, GLN111-U7, ARG58-C1, ARG58-G2, ARG109-U3, and ARG62-U6, which contribute to the stability of *raiZ* in State-2 (Figure S2-c in SI). The *raiZ* stayed attached to the

ProQ in this state throughout the rest of the simulation time. The RMSF analysis (Figure S3 in SI) reveals that in the raiz-ProQ complex, the amino acid residues 30-70, located in the binding site of State-1, have relatively higher RMSF values, indicating their possible role in reorganizing *raiZ* in the binding pocket.

### 2.3 Spontaneous Binding of *raiZ* with ProQ

Thus far, we have utilized a pre-docked ProQ-*raiZ* complex for our simulation studies. However, to get a comprehensive understanding of the binding process of our system, we must analyze our system from alternative perspectives as well. In this case, we conducted a spontaneous binding simulation (SBS) without any predetermined coordinates or path to explore other binding pockets of ProQ. We conducted multiple independent MD simulations, starting with different initial configurations of the ProQ-*raiZ* complex. For these simulations, we manually positioned the target RNA at varying distances (30-45 Å) and orientations relative to the ProQ. Multiple SBS results were presented in this study. The first outcome, depicted in Figure S4 (a-1), shows that the *raiZ* is positioned approximately 45 Å away from the protein. Initially, the *raiZ* was freely floating around; however, after a few nanoseconds, it began to approach the protein. This movement was primarily driven by long-range electrostatic interactions between positively charged amino acids and the negatively charged backbone of the *raiZ*. Eventually, the *raiZ* got attached to the lower end of the ProQ, as shown in Figure S4 (a-2).

In the second simulation, *raiZ* was positioned approximately 33 Å away from the ProQ on the upper side (Figure S4 (b-1) in SI). Initially, the *raiZ* attached to the front side of the ProQ is driven solely by electrostatic interactions, exploring other conformations over the protein surface. After some time, it started moving closer to the backside of the ProQ and ultimately got attached there in a relatively stable conformation compared to other conformations shown in Figure S4 (b-1).

### 2.4 ProQ-*raiZ* binding thermodynamics and mechanism

The equilibrium MD simulations including SBS provided pertinent atomistic insights into the binding interactions between the ProQ and *raiZ*. However, when delineating the RNA binding mechanism, equilibrium MD simulations find limitations due to the limited sampling time. We anticipate that *raiZ* binding to ProQ involves a much larger timescale, which standard brute-force MD simulations cannot sample. To circumvent this timescale limitation, we carried out On-the-fly Probability Enhanced Sampling (OPES) simulations^18^. We utilized the recently developed Deep-TDA method^21^ to design a collective variable (CV) based on 26 contact pairs between the ProQ and *raiZ* (Figure S5 in SI). In particular, we included many short and long-range contacts, including H-bonds, as descriptors for the DeepTDA method (see Methods section). Along with this Deep-TDA CV, we defined the backbone RMSD of *raiZ* as the second CV(s_2_). The use of Deep TDA along with a physical CV has been found effective in studying biomolecular recognition.

With these two CVs, s_1_ and s_2_, we carried out an OPES simulation to study the binding and unbinding mechanism of the ProQ-*raiZ* complex and explore possible metastable states, which are often inaccessible from equilibrium MD simulations due to the high energy barrier between the two metastable states. In the 2.5 μs OPES simulation, we observed a few back-and-forth transitions between the *raiZ*-bound state (−10) and unbounded state (10). (Figure S6 in SI) Using a reweighting method^19^, we calculated the free energy surface (FES) as a function of the Deep-TDA CV(s_1_) and another CV(s_3_) based on the coordination number of water (*j*=oxygen) surrounding an atom (1) at the center of *raiZ*, defined as

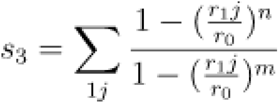

Here, *r*_1 *j*_ denotes the contact distance between the atom (*1)* of raiZ and oxygen atoms *(j)* of water, while *r*^0^ is a reference distance, set to 0.4 nm, that defines the distance range over which contact is considered significant; *n* and *m* are parameters of the switching function. The rationale for considering the third CV(s_3_) is that it could differentiate the basins well, but cannot efficiently sample them due to possible hysteresis.

In Figure 3, the basin (a) represents the conformation having the maximum number of contacts between ProQ and *raiZ* according to our Deep-TDA CV and CN, around 5, with water molecules around the binding pocket. This conformation is the same as State-1, which we initially conjectured as the binding pose of *raiZ* complexed with ProQ. Due to the high flexibility of the *raiZ* backbone and also the flexible loop region around the binding pocket of ProQ protein, *raiZ* explores many other metastable states having free energy values ranging from 0-80 kJ/mol, as shown in Figure 3e. By thoroughly analyzing all these metastable states (basins) on the FES, we found the possible pathway of *raiZ* binding to ProQ and subsequent folding. During unbinding, with the bias deposited in the basin (a), the *raiZ* moves to a more stable basin (b) in which it remains attached to the same concave surface but in a different orientation compared to the basin (a). Further deposition of bias in basin (b) leads to detachment of the RNA from the protein, and following basins (c) and (d), *raiZ* finally detaches completely from the ProQ surface and attains the unbound state. Similarly, during re-binding, it starts from basin (d) where positive amino acids attract the negatively charged *raiZ* and loosely bind the RNA. This is followed by the binding of *raiZ* to the lower side of ProQ, basins (c) and (d). Subsequently, *raiZ* starts to approach the binding pocket basin (b) in which positive amino acids ARG80, ARG58, ARG62, ARG109, and ARG100 mediate the binding process. Eventually, *raiZ* fits inside the binding pocket of ProQ and attains state (b), which is more stable than State-1 of the basin (a) obtained from docking. To characterize these basins further, we have carried out short ∼200-400 ns simulations starting from a configuration in each basin. From these trajectories, we calculated the two CV values and projected them on the FES. The trajectories sample the corresponding basin, indicating that the basins are intermediate states connecting the *raiZ-*bound and the unbound states. (Figure S8 in SI) To evaluate the convergence of the simulation, we conducted the subsequent tests: we examined the free energy difference (ΔG) as a function of simulation time, as illustrated in Figure S7. The plot indicates that the ΔG remains relatively constant after around 1 μs, implying that the free energy has almost converged.

**Figure 3:**
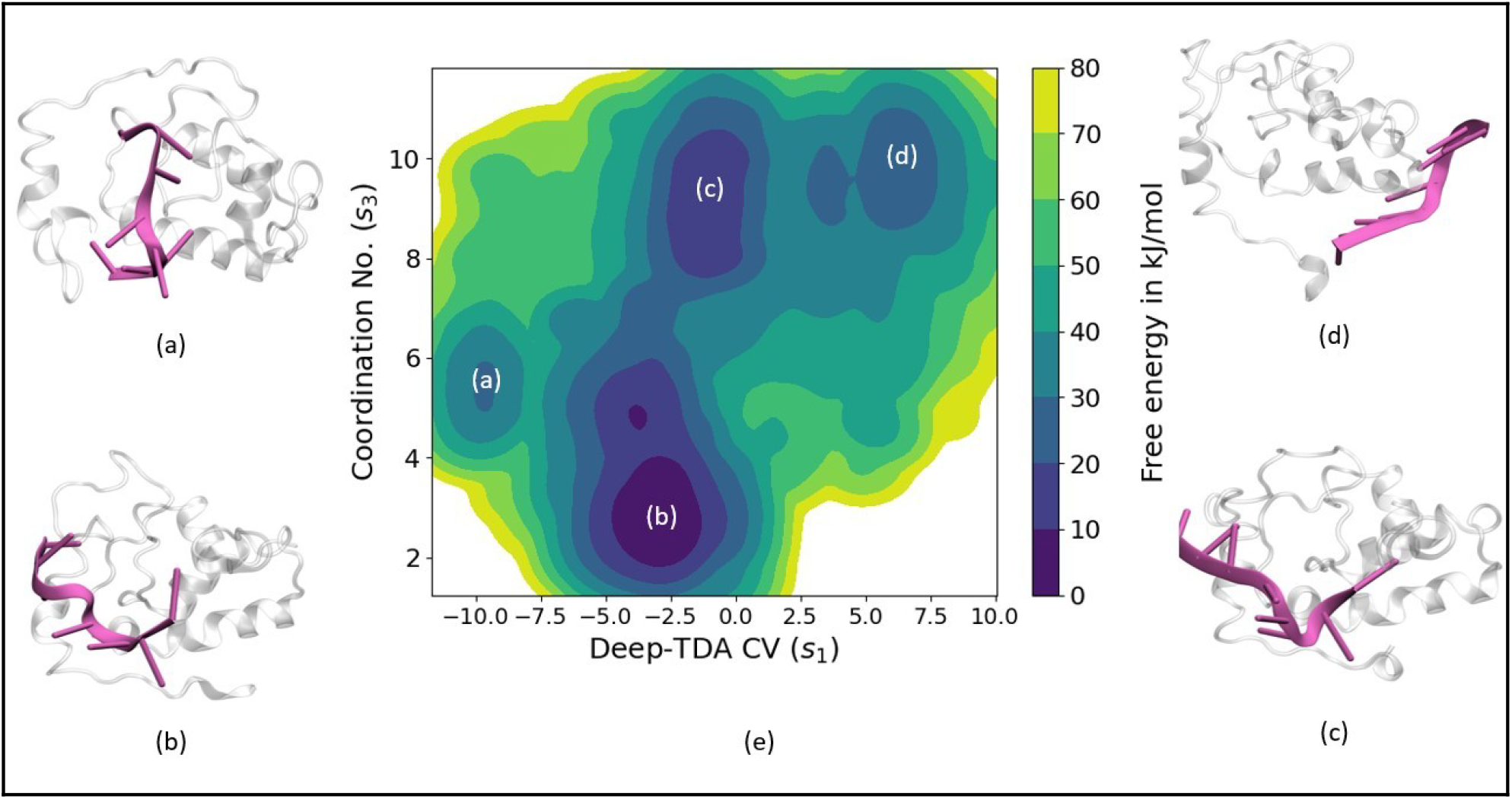
Free Energy Surface (FES) diagram with different metastable states - basin (a): initial bound state (State-1), basin (b): the deepest basin corresponding to the raiZ bound state in the same binding pocket but with a different orientation, basin (c): intermediate state similar to the basin (b) but having less contact with ProQ and having high CN represent more soluble with water also loosely bind to ProQ, basin (d): almost unbound state having very fewer contacts with ProQ and highly soluble with water as the higher value of CN, (e) the free energy surface (FES) as a function of s_1_ and s_3_ CVs.

Additionally, to elucidate the chaperone activity of ProQ, we computed the radius of gyration (R_g_) profiles of *raiZ* from the unbiased simulations initiated from each of the four basins (a), (b), (c), and (d) of the FES (Figure 3) and compared them with the R_g_ profile obtained from the simulation of a free *raiZ* in water (Figure S9 in SI). As evident from Figure S9, in water, *raiZ* prefers to stay in the extended conformation stabilized by water solvation, while when bound to ProQ, it gets partially folded, indicated by the reduced R_g_ values. In the most stable basin (b), raiZ has the most compact structure compared to the other basins. This analysis indicates the likely role of ProQ in RNA chaperone activity.

### 2.5. *In silico* mutations – the role of pocket residues

From the OPES simulation, we have found that in basin (a) (Figure 3a), ARG80 present inside the binding pocket stabilizes the *raiZ*. This residue has been found to play a crucial role in binding sRNA, experimentally proven by Pandey *et al*.^22^. To confirm the role of R80 in binding *raiZ* to ProQ, we replaced it with Alanine (ALA) and carried out a 500 ns long classical MD simulation. We plotted the distance between the Cα of ARG80 amino acid and U8 of the *raiZ* backbone atom to monitor the *raiZ’s* position with respect to the ProQ. The distance increases from 0.5 Å (in WT) to around 4.5 Å in mutants, indicating partial detachment of *raiZ* from the ProQ binding site (Figure S10 in SI). We repeated the simulations two more times to confirm the observation.

To further study the system, we analyzed the electrostatic potential surface of ProQ (Figure 4a), which shows a blue patch mainly consists of positively charged residues R69, R100, K101, and K107, which corresponds to the basin (b) in Figure 3b, the most stable binding confirmation we found from OPES simulation. The positively charged residues stabilize *raiZ* in the ProQ binding pocket. We carried out *in silico* mutations to confirm their role in *raiZ* binding. Single point mutation of each of the three residues to ALA did not have much effect on *raiZ* binding, with no significant detachment of *raiZ* from ProQ (Figure S11 in SI). Replacing LYS101 with ALA leads to partial detachment of *raiZ*. (Figure S11 in SI). This is not quite unexpected since a single point mutation of a positively charged residue is insufficient to remove the electrostatic interaction between the negatively charged *raiZ* and positively charged blue patch of ProQ. So we prepared a triple mutant by replacing all three amino acids ARG69, ARG100, and LYS101 together with ALA. We calculated the distance between C-alpha of ALA100 of ProQ and C1 of *raiZ* to monitor *raiZ* unbinding (Figure S12 in SI). We observed the distance increase sharply compared to WT (0.5 Å) to around 4 Å in the triple mutant. Repeat simulation also provided similar outcomes. This suggests that it is not a single amino acid that is crucial for *raiZ* binding with ProQ, but the complete blue patch (Figure 4a) of positively charged amino acids ARG69, ARG100, and LYS101 holds *raiZ* on the binding site.

**Figure 4.**
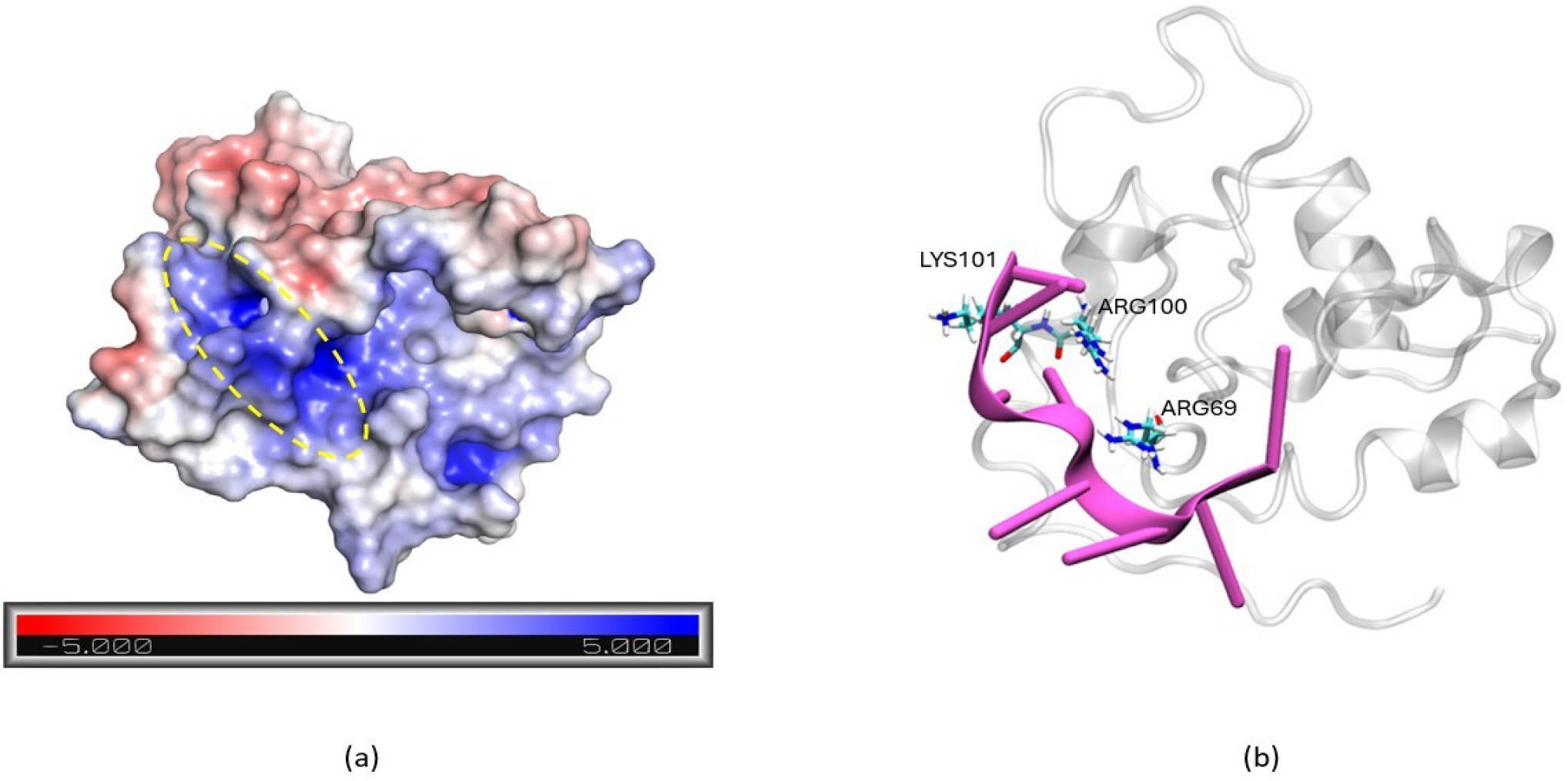
(a) Electrostatic potential surface for ProQ: the positively charged region shown in blue is highlighted by a yellow circle, and (b) shows amino acids forming a blue patch in ProQ for binding *raiZ*.

## Conclusion

In summary, we have investigated the specific action of ProQ in small RNA *raiZ* binding and subsequent folding. Utilizing long microsecond timescale molecular dynamics (MD) simulations, we found the conformational preferences of *raiZ* RNA on the ProQ’s concave-shaped binding pocket and identified positively charged amino acids such as ARG, LYS, and HIS playing a crucial role in holding the RNA inside the binding site and aiding in its folding. In addition, we utilized OPES simulations to explore the mechanism of *raiZ* binding to ProQ and calculate the free energy landscape, and found that basin (b) is the most stable one. Initially, the *raiZ* interacts with ProQ’s surface through its positively charged residues; as conformational fluctuations continue, the *raiZ* gradually approaches the binding site where ProQ folds *raiZ* RNA. Moreover, the low radius of gyration of *raiZ* in the bound state basin (b) supports the RNA Chaperone activity of ProQ. *In-silico* mutational studies revealed that it is not a single amino acid, but rather a positively charged patch on ProQ, comprised of ARG69, ARG100, and LYS101 plays a crucial role in binding *raiZ*. These findings substantially improve our knowledge of ProQ’s role as an RNA chaperone and its interaction with small RNAs like *raiZ*. The knowledge gathered from this research serves as a valuable foundation for future investigations conducted in *in vitro* and *in vivo* studies. This positions the ProQ-*raiZ* complex as a promising target for the development of new antibiotics.

## Methodology

### Dataset

We extracted the NMR structure of the N-terminal domain of the Escherichia Coli ProQ RNA binding protein from the RCSB^23^ protein data bank (PDB ID: 5NB9; Grecia M. Gonzalez et. al.).^24^ Out of 17 models present in the ProQ PDB file, the first model structure was used for this investigation, since it was close to the average structure and also had the lowest energy value among 17 models. For generating the sRNA (*raiZ*) 3D structure file, the sequence was downloaded from the web database EcoCyc^25,26^ (*raiZ* accession ID: GO-17078). Further, the RNAfold^27^ and RNA composer tool^28,29^ were used for developing the coordinates in PDB format.

### Molecular Docking studies

Molecular docking of the N-terminal domain of the Escherichia Coli ProQ protein, along with sRNA *raiZ* was performed using the AutoDock Vina software^20^. For docking purposes, only 8 nucleoside long sequences of the putative binding site (5’-CGUUUUUU-3’), which is present on the 3’ end of the *raiZ*, have been used. The binding affinity was computed for this ProQ-raiZ complex. For the grid box, the XYZ coordinates were set at -1.242, -7.319, and 4.461, respectively, with a grid spacing of 1 Å. The dimensions of the box were set at 80×80×80 with an exhaustiveness value equal to 8 and an energy range of 4. From docking results, we got 9 different binding conformations having different binding affinities, out of which the first conformation was selected having the highest binding affinity equal to -7.5 kcal/mol. This ProQ-raiZ complex has been used for further molecular dynamics studies.

### Molecular dynamics simulations

The structure obtained from the molecular docking result was utilized for molecular dynamics simulation. We carried out three sets of simulations - (i) ProQ in water, (ii) putative binding site (5’-CGUUUUUU-3’) of *raiZ* in water, and (iii) ProQ-*raiZ* docked complex in water using GROMACS v 2021.4.^30^ The AMBER99SB-ILDN force field^31^ was used for ProQ. While, for *raiZ*, the OL3^32^ nucleic acid force field was used for generating the interaction potentials both in water as well as in complex. In all the systems, the simulation box was solvated using the SPCE water model^33^, and ions were added to neutralize the system.^34^ Further, the steepest descent algorithm was used to remove the unwanted contacts and bring the system to a minimum energy. Lastly, all three systems were equilibrated, using the v-rescale^35^ thermostat and the Parrinello-Rahman barostat^36^ to a target temperature of 300 K and 1 bar pressure, respectively. The production MD was performed using the leap-frog algorithm and equations of motions were integrated after every 2 fs.

### Enhanced Sampling Method

We employed the recently developed enhanced sampling method, the On-the-fly Probability Enhanced Sampling (OPES)^18,37^ to sample binding and unbinding events and explore a range of possible conformations of the system. The Deep-TDA CV (s_1_) along with the raiZ RMSD CV (s_2_), were used in the OPES simulations to explore the range of possible conformations of the system. To train the Deep-TDA CV, we took 26 contact pair distances between the Cα atoms of amino acids of the ProQ protein and the backbone atoms of *raiZ* RNA. These contacts were chosen from the equilibrium simulation trajectories. The training was done for two distinct bound states and one unbound state. Afterward, utilizing these descriptors, the Deep-TDA CV was constructed. The Pytorch40^38^ library was utilized for training the model, and subsequently, the trained model from Pytorch was imported into PLUMED utilizing the interface built by Bonati *et al*.^39^. The GROMACS-2021.4^30^ patched with PLUMED2.9^40^ was utilized to conduct the enhanced sampling (ES) simulations.

## Acknowledgments

The authors acknowledge IIT Delhi for the research funding and all necessary facilities. SS and NK thank IITD for their Ph.D. fellowships. T.K. thanks the IITD for the seed grant. The authors extend their appreciation to the IITD HPC for the supercomputing facility (DST-FIST).

## Supporting Information

The supporting information contains the binding affinity of different binding modes obtained from docking of ProQ and *raiZ* using AutoDock, RMSD, and RMSF plots, spontaneous binding simulation results, CV profile, FES with projected CV values from unbiased simulations run at each basin of the FES, Rg plot showing ProQ’s chaperon activity.

## Data Availability Statement

The input files required to run the simulations and results data can be found in the GitHub repository https://github.com/ShilpiSinghIITD/ProQ_raiZ.git

