## Supplemental Information for "Understanding RNA Chaperone Activity of ProQ Protein"

| Binding mode | Binding affinity (kcal/mol) |
|---|---|
| 1 | -10.8 |
| 2 | -10.5 |
| 3 | -10.4 |
| 4 | -10.4 |
| 5 | -10.1 |
| 6 | -10.1 |
| 7 | -10.1 |
| 8 | -10.1 |
| 9 | -10.1 |

**Table S1:** Binding affinity of different binding modes obtained from docking of ProQ and *raiz* using AutoDock

### 1. RMSD and RMSF analysis

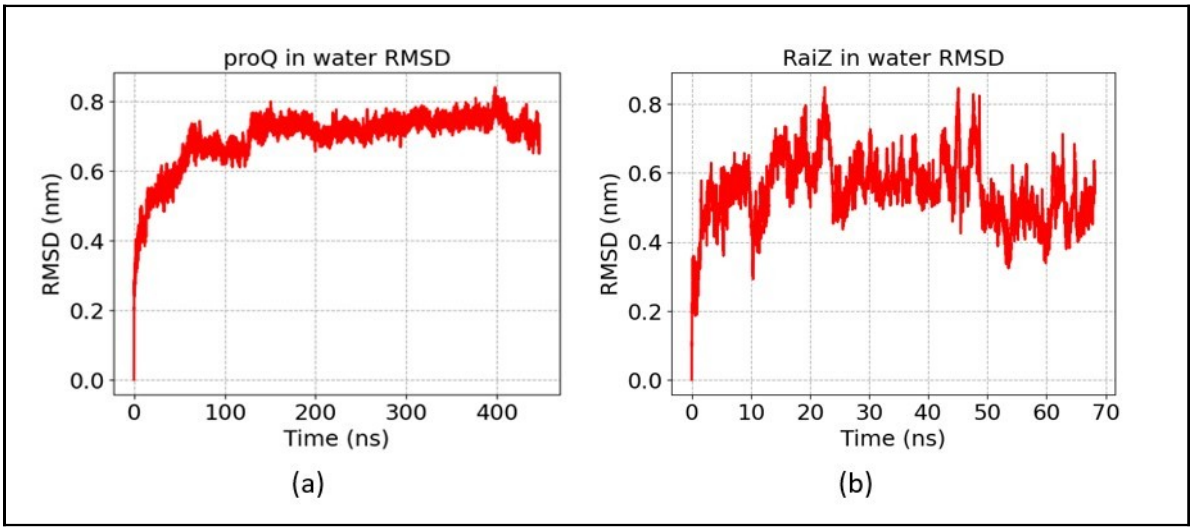


**Figure S1:** (a) RMSD vs time plot for ProQ in water and (b) RMSD vs time plot for *raiZ* in Water

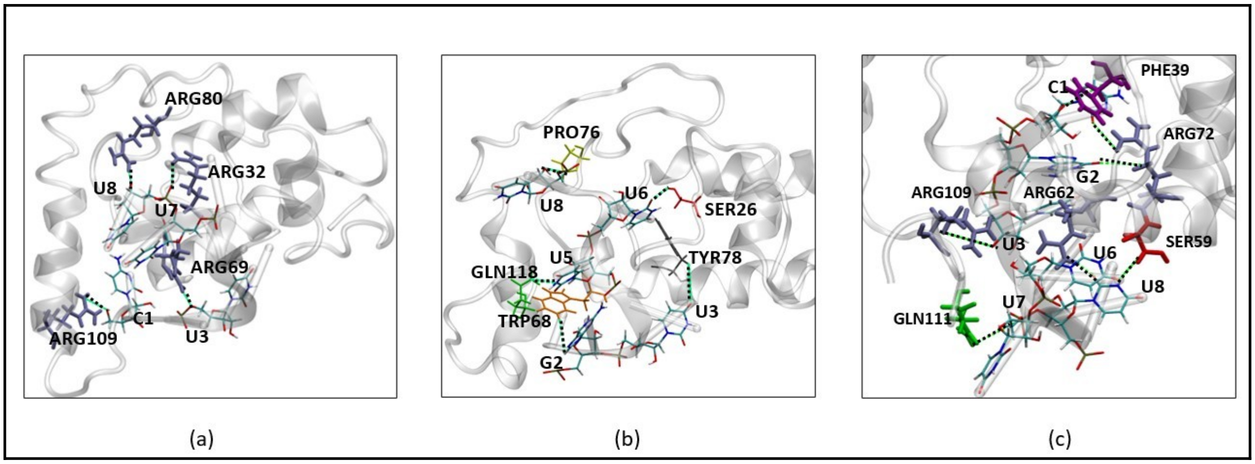


**Figure S2:** (a) Electrostatic interactions occurred between positively charged ARG of ProQ and *raiZ* in State-1, (b) residues showing H-bonding between ProQ and *raiZ* in State-1 and (c) Electrostatic interactions occurred between positively charged amino acids of ProQ and negatively charged residues of *raiZ* in State-2

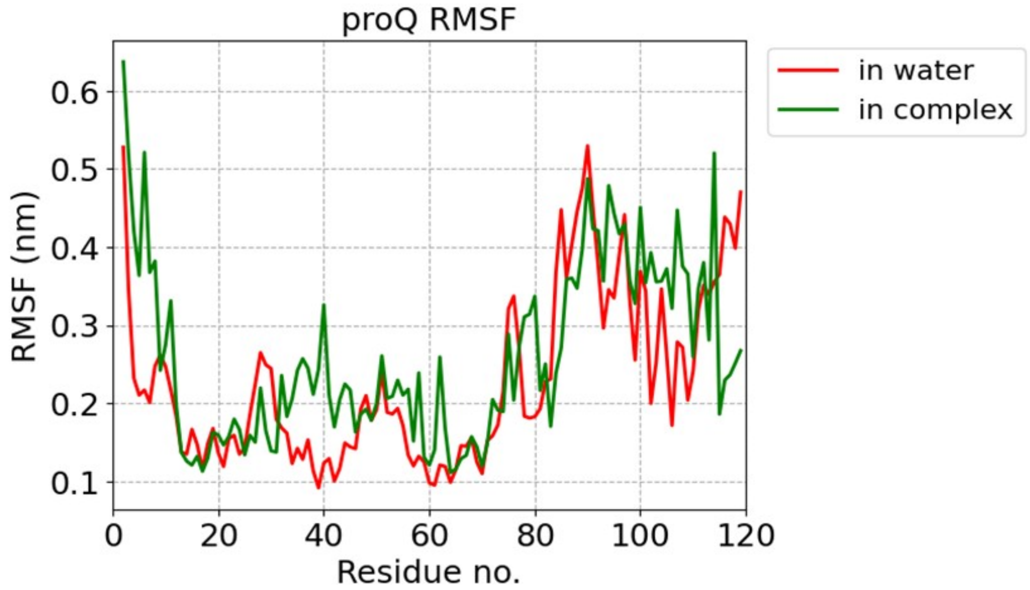


**Figure S3:** Root mean square fluctuation Plot

### 2. Spontaneous Binding Simulation (SBS)

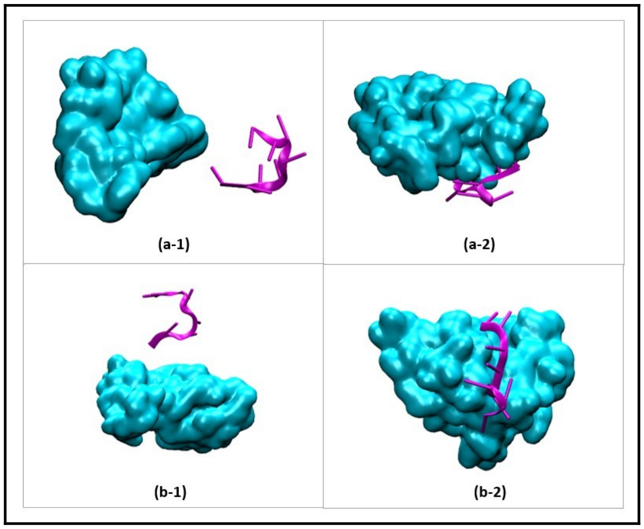


**Figure S4:** Binding conformation of *raiZ* to protein during Spontaneous Binding Simulation (SBS)

### 3. Thermodynamics study

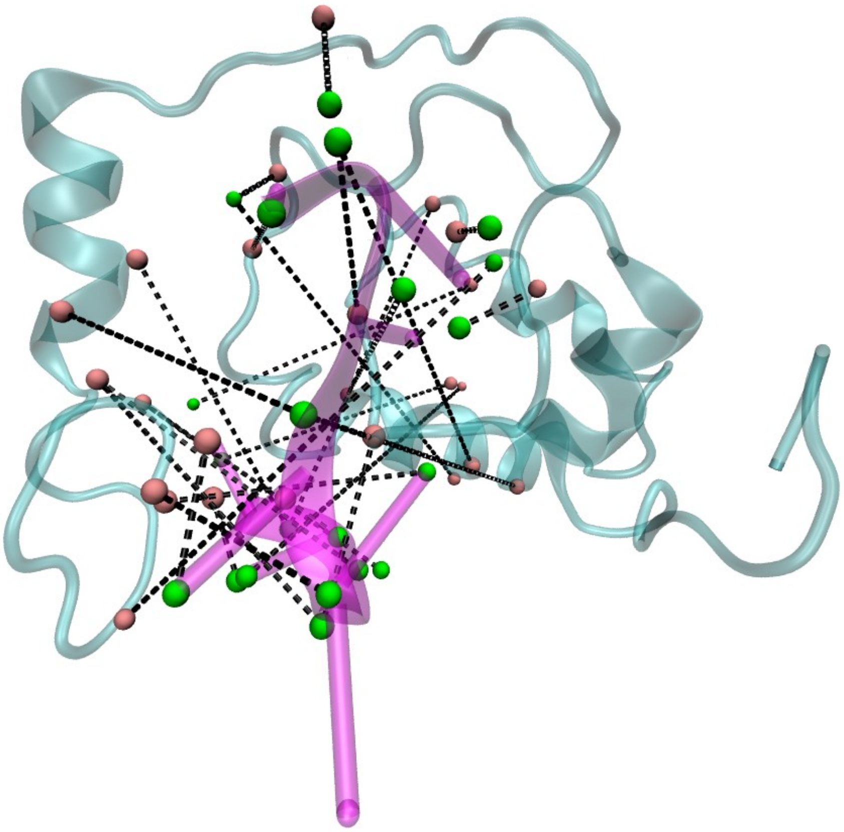

**Figure S5:** Long and short Contact pairs between ProQ-*raiZ* (balls in light pink represents ProQ protein atom and green balls represents *raiZ* rna atom in contact pair )

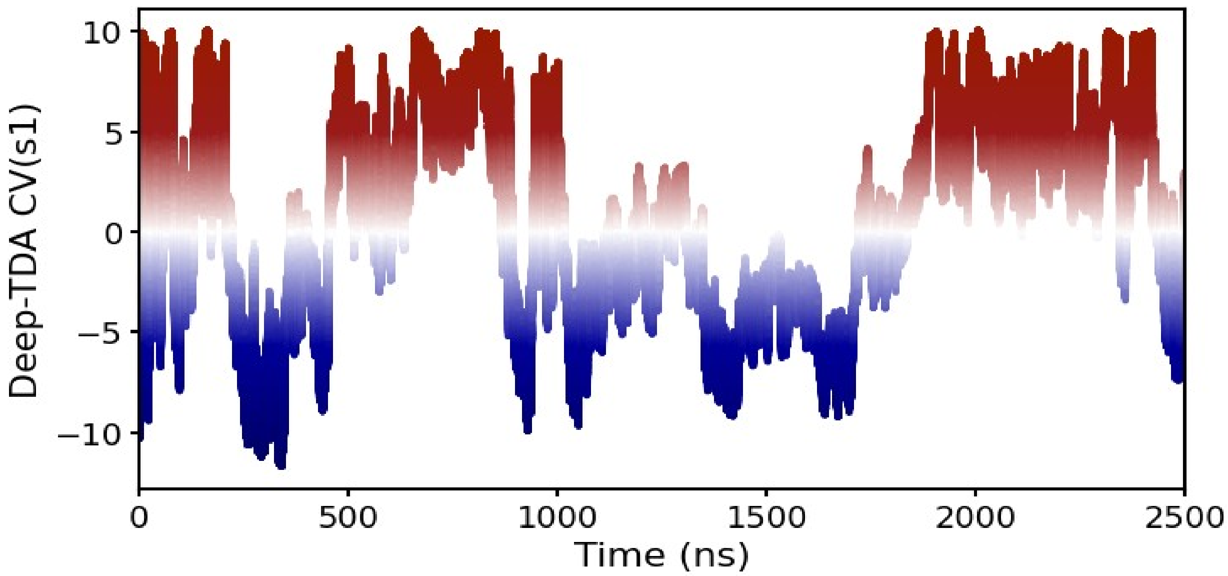


**Figure S6:** Back-and-forth transitions between the RNA bound state (-10) and unbounded state (10) vs time.

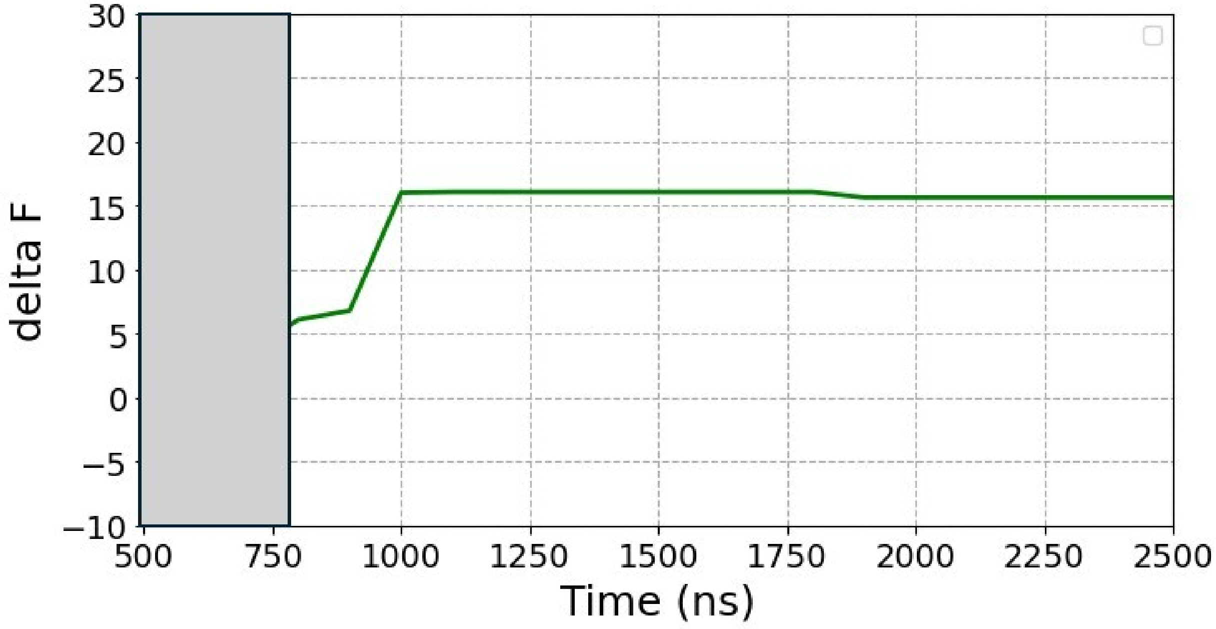


**Figure S7:** deltaF vs time which shows change in free energy of running FES depicts the convergence

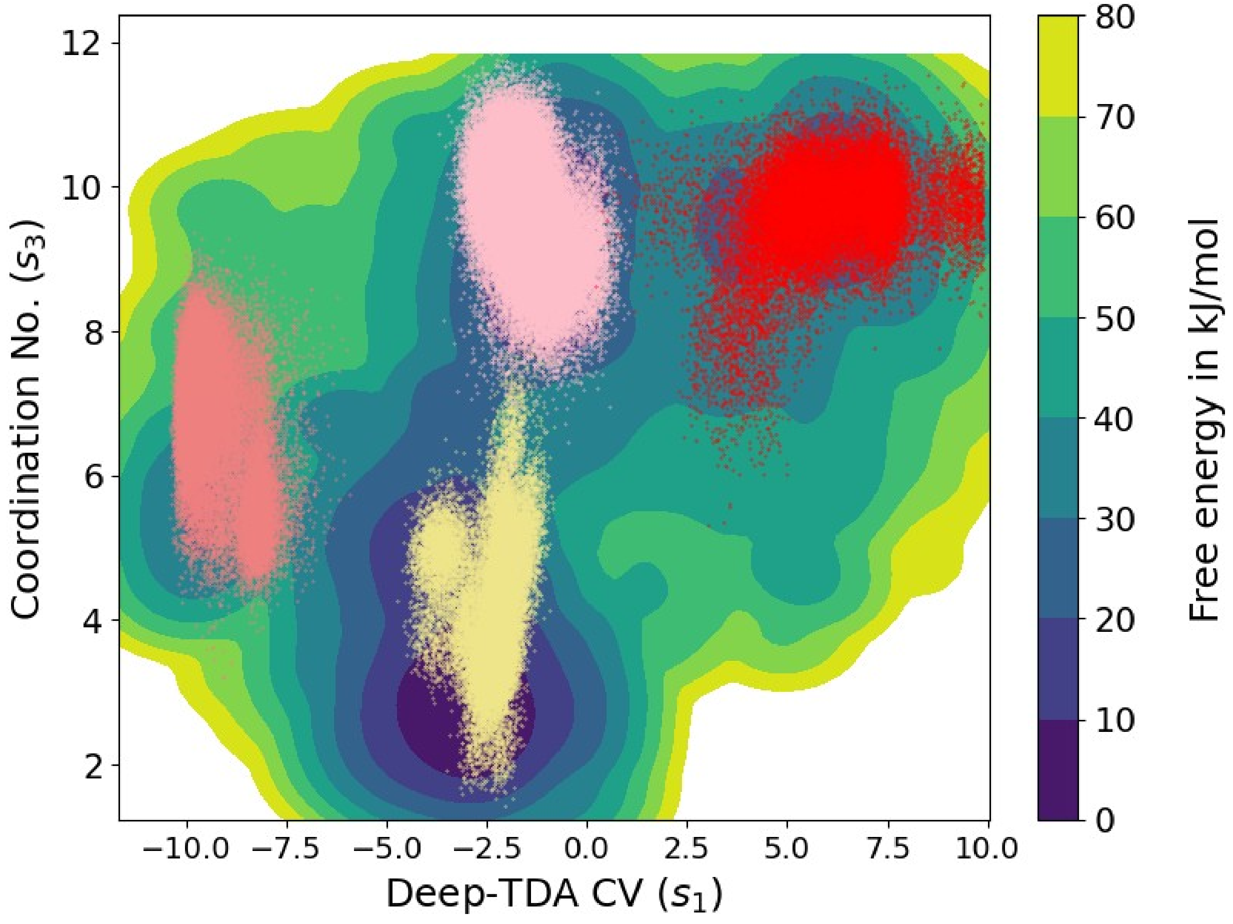


**Figure S8:** The free energy surface (FES) in kJ/mol is calculated as a function of the CN between the *raiZ* backbone and surrounded water atoms, and Deep-TDA CV is obtained for contacts between ProQ-*raiZ*. The CVs data obtained from four equilibrium simulation trajectories are projected onto the FES using the following color: light coral corresponds to the transition state near basin (a), yellow represents the state near basin (b), light pink corresponds to the state in basin (c) and red shows transition state near basin (d).

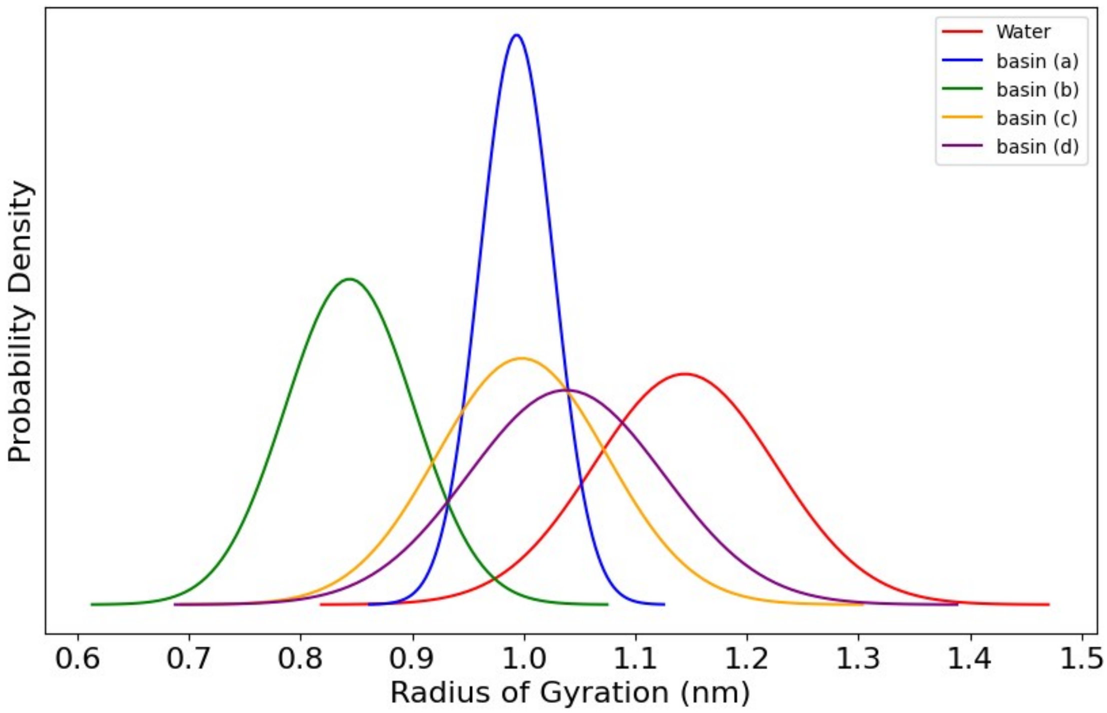


**Figure S9:** Radius of gyration (Rg) vs probability density graph, red Gaussian shows *raiZ* in water without ProQ having high Rg values, green Gaussian represents lowest Rg values represents folding of *raiZ* which supports chaperone activity of ProQ.

### 4. In-silico Mutations

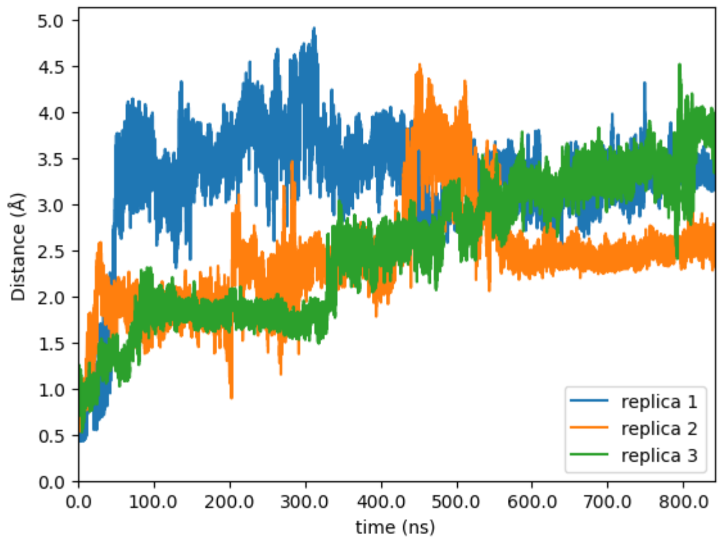


**Figure S10:** Distance plotted between C-alpha of ALA80 of ProQ and U8 of *raiZ* for each replica

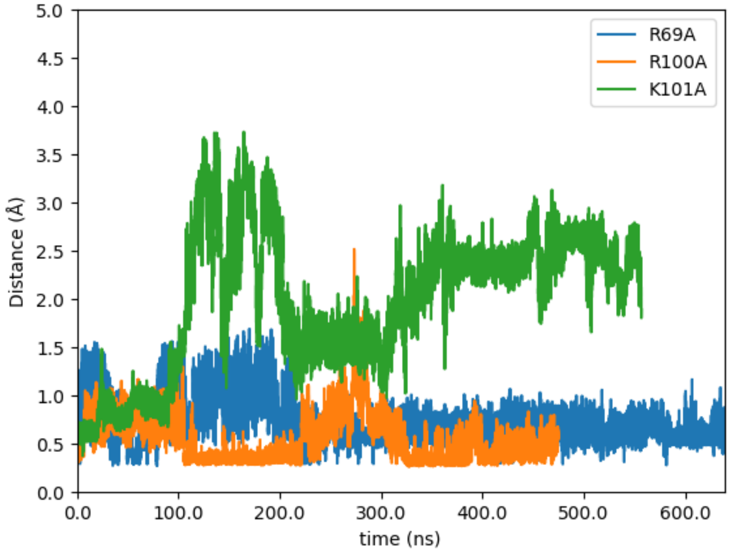


**Figure S11:** Distance plotted between C-alpha of ALA69 of ProQ and C1of *raiZ* (blue) , C-alpha of ALA100 of ProQ and C1 of *raiZ* (orange), and C-alpha of ALA101 of ProQ and C1 of *raiZ* (green)

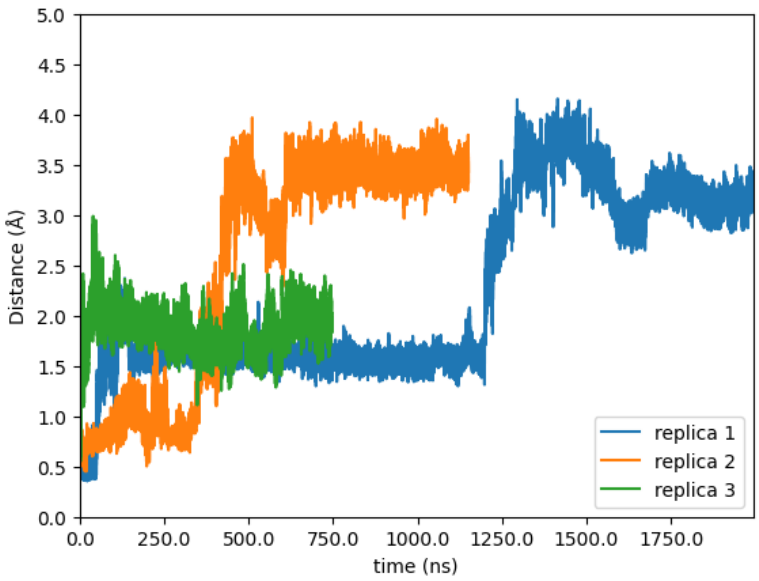


**Figure S12:** Distance plotted between C-alpha of ALA100 of ProQ and C1 of *raiZ* for each replica
